# PARP Inhibition Impedes the Maturation of Nascent DNA Strands During DNA Replication

**DOI:** 10.1101/2021.07.03.450982

**Authors:** Alina Vaitsiankova, Kamila Burdova, Hana Hanzlikova, Keith W Caldecott

## Abstract

PARP1 is implicated in the detection and repair of unligated Okazaki fragment intermediates, highlighting these structures as a potential source of genome breakage induced by PARP inhibition. In agreement with this, we show here that PARP1 activity is greatly elevated in chicken and human S phase cells in which FEN1 nuclease is genetically deleted, and that PARP activity is highest tens of kilobases behind DNA replication forks. Importantly, PARP inhibitor reduces the integrity of nascent DNA strands in both wild type chicken and human cells during DNA replication, and does so in *FEN1^−/−^* cells to an even greater extent that can be detected as post-replicative single-strand gaps within individual DNA fibres. Collectively, these data show that PARP inhibitors impede the maturation of Okazaki fragments in nascent DNA, implicating these canonical DNA replication intermediates in the cytotoxicity of these compounds.

## Introduction

Poly(ADP-ribose) polymerases (PARPs) are a superfamily of enzymes that utilise NAD+ to modify themselves and other proteins with mono- or poly(ADP-ribose) ^1,2^. The archetypal PARP enzyme is PARP1 which, along with PARP2 and PARP3, is activated by DNA breaks and regulates the cellular DNA damage response ^3–5^. Poly(ADP-ribose) is a highly dynamic and transient signal that is rapidly degraded by poly(ADP-ribose) glycohydrolase (PARG) ^6–10^. DNA damage-stimulated PARPs bind to and are activated by a variety of DNA substrates, of which DNA single- and double-strand breaks (SSBs and DSBs) are the best characterised. At SSBs, PARP1 and PARP2 fulfil a variety of roles, depending on the nature and source of the break, including the regulation of chromatin compaction and the recruitment of DNA repair proteins ^5,11^.

In addition to DNA breaks arising stochastically across the genome, PARP1 and PARP2 are also involved in the detection and processing of various DNA replication intermediates ^12^. Indeed, S phase is the primary source of poly(ADP-ribosylation) in unperturbed proliferating cells ^13^. For example, PARP1 may detect and signal the presence of paused, reversed, and/or collapsed DNA replication forks ^14,15^. A likely role for PARP1 and/or PARP2 at collapsed forks is to suppress binding by Ku and 53BP1, which otherwise can trigger ‘toxic’ non-homologous end-joining (NHEJ) ^16–18^. In addition, PARP activity may promote homologous recombination (HR)-mediated resetting and/or repair of reversed or collapsed forks, by regulating the recruitment and/or activity of MRE11 nuclease ^14,19^. PARP1 can also regulate the longevity of reversed forks by inhibiting RECQ1, a helicase that can reset reversed forks independently of RAD51-mediated HR ^20,21^.

Recently, we implicated PARP1 also in the detection of unligated Okazaki fragments ^13^. The synthesis of Okazaki fragments is initiated by DNA polymerase α-primase complex (POLα), which generates short RNA primers that are extended by POLα for 10-20 deoxyribonucleotides followed by DNA polymerase δ (POLδ) for a further ~200 deoxyribonucleotides, until the 5’-terminus of the downstream Okazaki fragment is encountered ^22–26^. The junctions between adjacent fragments are then processed and ligated by flap endonuclease-1 (FEN1) and DNA ligase I (LIG1), respectively, though other nucleases may be involved ^22–27^. Whilst this canonical pathway for the maturation of Okazaki fragments is very efficient, it has been estimated from biochemical experiments that ~15-30% of human POLδ molecules disengage before reaching a downstream Okazaki fragment, even in the presence of the PCNA processivity factor ^28^. Given that each human S phase entails the formation of 30-50 million Okazaki fragments, failure of the canonical pathway to ligate even just 0.1% of Okazaki fragment intermediates would result in 30-50 thousand SSBs and/or single-strand gaps each S phase. PARP1-dependent signalling and repair may thus help ensuring the integrity of nascent DNA strands during normal DNA replication. Consistent with this idea, SSB repair proteins recruited at DNA breaks by PARP1 such as X-ray repair cross-complementing protein 1 (XRCC1) and DNA ligase III (LIG3) have been associated with Okazaki fragment maturation ^13,29–31^.

Inhibitors of PARP are clinically approved drugs for the treatment of cancer cells in which HR-mediated repair is defective, on the basis of their extreme toxicity in such cells ^32–34^. A critical mechanistic aspect of such inhibitors is their ability to ‘trap’ PARP enzymes on their endogenous DNA substrates, which in the absence of efficient HR-mediated repair leads to cell death ^34,35^. However, the critical DNA substrates on which PARP inhibitors trap PARP enzymes, and the impact of this trapping on normal DNA metabolism during DNA replication are poorly understood. Here, we have addressed this question and we find that PARP inhibitors impede the maturation of nascent DNA strands during DNA replication, and we implicate unligated intermediates of Okazaki fragment processing as a major source of cytotoxic PARP1 trapping.

## Results

To test our hypothesis that PARP1 becomes ‘trapped’ on unligated Okazaki fragments, we employed wild type chicken DT40 cells and their derivatives in which FEN1 was deleted by gene targeting ^31^. *FEN1*^−/−^ DT40 cells are viable and proliferate, albeit with slightly increased doubling time that likely reflects increased cell death (Extended Data Fig.1A), suggesting that alternative pathways for Okazaki fragment maturation operate in these cells. To examine whether PARP1 contributes to these pathways we measured the steady-state level of ADP-ribosylation in wild type and *FEN1^−/−^* DT40 cells. Consistent with this idea, a short incubation with PARG inhibitor to preserve nascent poly(ADP-ribose) revealed elevated levels of ADP-ribosylation in *FEN1^−/−^* DT40 cells, specifically in S phase, when compared to wild type DT40 cells (Fig.1A & Extended Data Fig.1B). S phase ADP-ribosylation was also increased in *FEN1^−/−^* DT40 cells in the absence of PARG inhibitor, albeit this increase fell short of statistical significance (Fig.1A). The amount of PARP1 present in the detergent-insoluble fraction of *FEN1^−/−^* DT40 cells was also elevated when compared to wild type cells, and was increased further by incubation with PARP inhibitor, consistent with the ‘trapping’ of PARP1 on unligated Okazaki fragments (Fig.1B). Notably, *FEN1^−/−^* DT40 cells were more sensitive to PARP inhibitor than were *XRCC3^−/−^* DT40 cells that lack efficient HR, which is the archetypal determinant of cellular sensitivity to PARP inhibitors, suggesting that PARP1 trapping on Okazaki fragments is a highly toxic event (Fig.1C).

**Figure 1.**
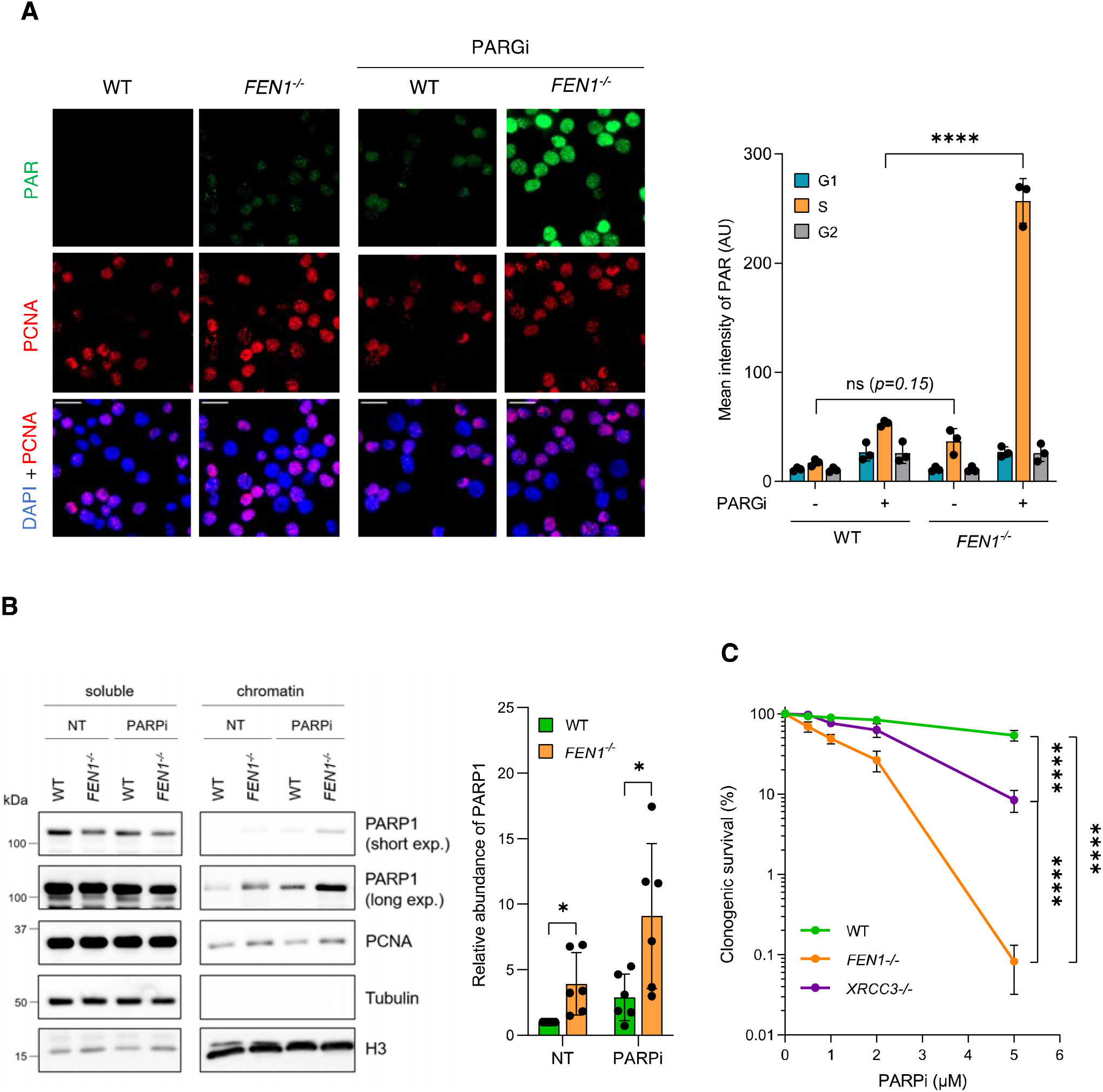
Increased PARP1 activity and sensitivity to PARP inhibitor in *FEN1^−/−^* DT40 cells. **(A)** Representative images (*left*) and ScanR quantification (*right*) of poly(ADP-ribose) fluorescence (PAR) (MABE1031) in wild type (WT) and *FEN1^−/−^* DT40 cells incubated for 30 min with DMSO vehicle (−) or with 10 μM PARG inhibitor (PARGi) to prevent poly(ADP-ribose) degradation. S phase cells were distinguished by PCNA staining and G1/G2 phase cells by DAPI intensity. Data represent the average (±SD) mean PAR fluorescence in arbitrary units (AU) from 3 independent experiments with individual data points plotted. Scale bars, 20 μm. Statistical significance was assessed by 1-way ANOVA with post-hoc Sidak’s multiple comparisons test. Galleries of representative cells from ScanR analysis are shown in Figure S1B. **(B)** Western blots (*left*) of PARP1, PCNA, tubulin, and histone H3 (H3) in soluble (detergent extracted) and chromatin-containing fractions from wild type (WT) and *FEN1^−/−^* DT40 cells. Cells were incubated for 30 min with DMSO vehicle (NT) or 10 μM PARP inhibitor (PARPi; Olaparib) prior to cell fractionation. For quantification of PARP1 levels in chromatin (*right*), PARP1 protein levels in the detergent-insoluble material were first normalised to ponceau S-stained histone levels and then expressed relative to the PARP1 level in untreated wild type chromatin. Data represent the mean (±SD) of 6 independent experiments with individual data points plotted. Statistical significance was assessed by Student’s paired t test. **(C)** Clonogenic survival assays for wild type (WT), *FEN1^−/−^*, and *XRCC3^−/−^* DT40 cells grown with the indicated concentrations of PARPi (Olaparib). Data represent the mean (±SD) of 4 independent experiments. Statistical significance was assessed by 2-way ANOVA with post-hoc Tukey’s multiple comparisons test. For all panels; ns, not significant; *p< 0.05; **p<0.01; ****p<0.0001.

To examine directly whether PARP inhibitors might block the maturation of Okazaki fragments we measured the integrity of genomic DNA in wild type and *FEN1^−/−^* DT40 cells using alkaline comet assays. The level of endogenous DNA breaks was ~2-fold higher in *FEN1^−/−^* cells than in wild type cells, as measured by their comet tail moments (an arbitrary measure of DNA strand breaks), and this difference was increased a further ~2-fold by incubation with PARP inhibitor (Fig.2A & Extended Data Fig.2A). To confirm that the elevated DNA breaks in *FEN1^−/−^* cells were associated with DNA replication we measured the integrity of genomic DNA specifically in S phase. Indeed, most of the DNA breaks detected by alkaline comet assays were in S phase in both wild type and *FEN1^−/−^* cells (Fig.2B & Extended Data Fig.2B). Once again, the level of endogenous DNA breaks was ~2-fold higher in *FEN1^−/−^* cells than in wild type cells, and was elevated a further ~2-fold by PARP inhibitor (Fig.2B & Extended Data Fig.2B). The DNA breaks detected in S phase cells were not an artifact of labelling cells with BrdU, because omission of this nucleotide from experiments had no impact on the results of our alkaline comet assays (Extended Data Fig.2C).

**Figure 2.**
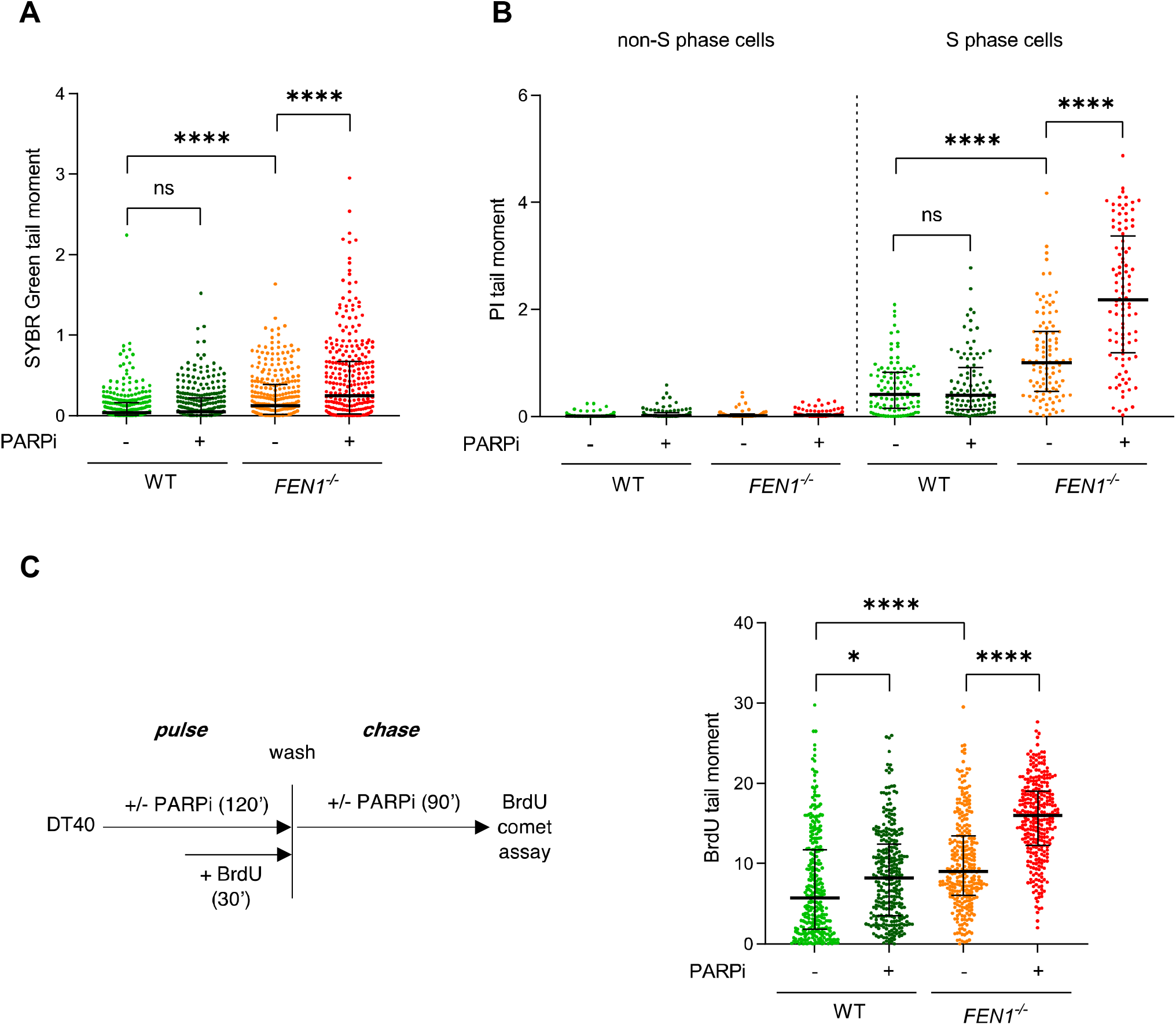
PARP inhibitor impedes the maturation of large/late nascent DNA strands in wild type and *FEN1^−/−^* chicken DT40 cells. **(A)** DNA strand breaks were quantified by alkaline comet assays in wild type (WT) and *FEN1^−/−^* DT40 cells following a 2 h incubation with DMSO vehicle (−) or 10 μM PARPi (KU 0058948). Genomic DNA was scored for comet tail moments by staining with SYBR Green. For each sample, scatter plots are the comet tail moments (an arbitrary unit of DNA strand breakage) of 300 cells combined from 3 independent experiments (100 cells per sample per experiment) and the bars represent the median and interquartile range. The individual data sets from the three experimental repeats are plotted in Figure S2A. Statistical significance was determined by 2-way ANOVA with Tukey’s post hoc multiple comparisons test, with individual tail moments within an experiment treated as technical replicates and with experimental matching/blocking by experimental repeat. **(B)** DNA strand breaks in total genomic DNA were quantified by alkaline comet assays in S phase and non-S phase cells following incubation or not with 10 μM PARPi (Olaparib) for 2 h. S phase cells were identified by labelling with BrdU for the final 45 min. Genomic DNA was scored for comet tail moments by staining with propidium iodide (PI). For each sample, scatter plots are the comet tail moments of 100 cells combined from 2 independent experiments (50 cells per sample per experiment) and the bars represent the median and interquartile range. The individual data sets from the two experimental repeats are plotted in Figure S2B, and statistical analysis was conducted as in panel A. **(C)** DNA breaks in nascent DNA strands quantified in DT40 cells by anti-BrdU alkaline comet assays after pulse labelling (30 min) with BrdU followed by a subsequent chase (90 min), as indicated in the schematic (*left*). Where indicated, 10 μM PARPi (Olaparib) was present in the media. Alkaline comet tail moments were quantified in BrdU-labelled nascent single-strands by staining with anti-BrdU antibodies (“BrdU comet tail moment”). For each sample, scatter plots show BrdU comet tail moments from 300 cells combined from 3 independent experiments (100 cells per sample per experiment) and bars are the median and interquartile range. The individual data sets from the three experimental repeats are plotted separately in Figure S2D, and statistical analysis was conducted as in panel A. For all panels; ns, not significant; *p< 0.05; ****p<0.0001.

To explore further whether the elevated DNA breaks in *FEN1^−/−^* cells reflected an impact of PARP1 trapping on the maturation of Okazaki fragments, we measured the integrity of nascent DNA strands following BrdU pulse-labelling and a 90 min chase (Fig.2C, *left*). Since DNA strands are separated in alkaline comet assays the quantification of tail moments using anti-BrdU antibodies can measure the integrity specifically of nascent DNA strands. Notably, nascent strand integrity was significantly reduced in *FEN1^−/−^* cells when compared to wild type DT40 cells, following BrdU pulse labelling and a 90 min chase, consistent with a reduced rate of nascent strand maturation in the mutant cells (Fig.2C & Extended Data Fig.2D). More importantly, PARP inhibitor reduced the integrity of nascent DNA strands in wild type DT40 cells, and did so to an even greater extent in *FEN1^−/−^* cells (Fig.2C & Extended Data Fig.2D).

Because BrdU comet assays are sensitive only to large nascent DNA fragments of 500 kb or more (Extended Data Fig.2E), we wanted to examine the impact of PARP inhibitor on smaller nascent DNA strands, reflecting early DNA replication intermediates. To do this, we employed alkaline agarose gels, in which the distribution of DNA fragments of <10 kb can be resolved (Fig.3A). An increased fraction of nascent DNA was present as fragments of 10 kb or less in *FEN1^−/−^* DT40 cells, when compared to wild type cells, following a 10 min pulse label with [^3^H]-thymidine, and remained so throughout a subsequent 20 min chase (Fig.3B). Moreover, although PARP inhibitor did not measurably impact on the amount of nascent DNA present as fragments of <10 kb in wild type DT40 cells, it had a significant impact on the amount of these fragments in *FEN1^−/−^* cells, increasing their prevalence during both pulse labelling and the subsequent chase (Fig.3B). Collectively, these data indicate that PARP inhibitor impedes the maturation of nascent replication intermediates in DT40 cells, and that this impact is particularly pronounced if canonical Okazaki fragment processing is perturbed.

**Figure 3.**
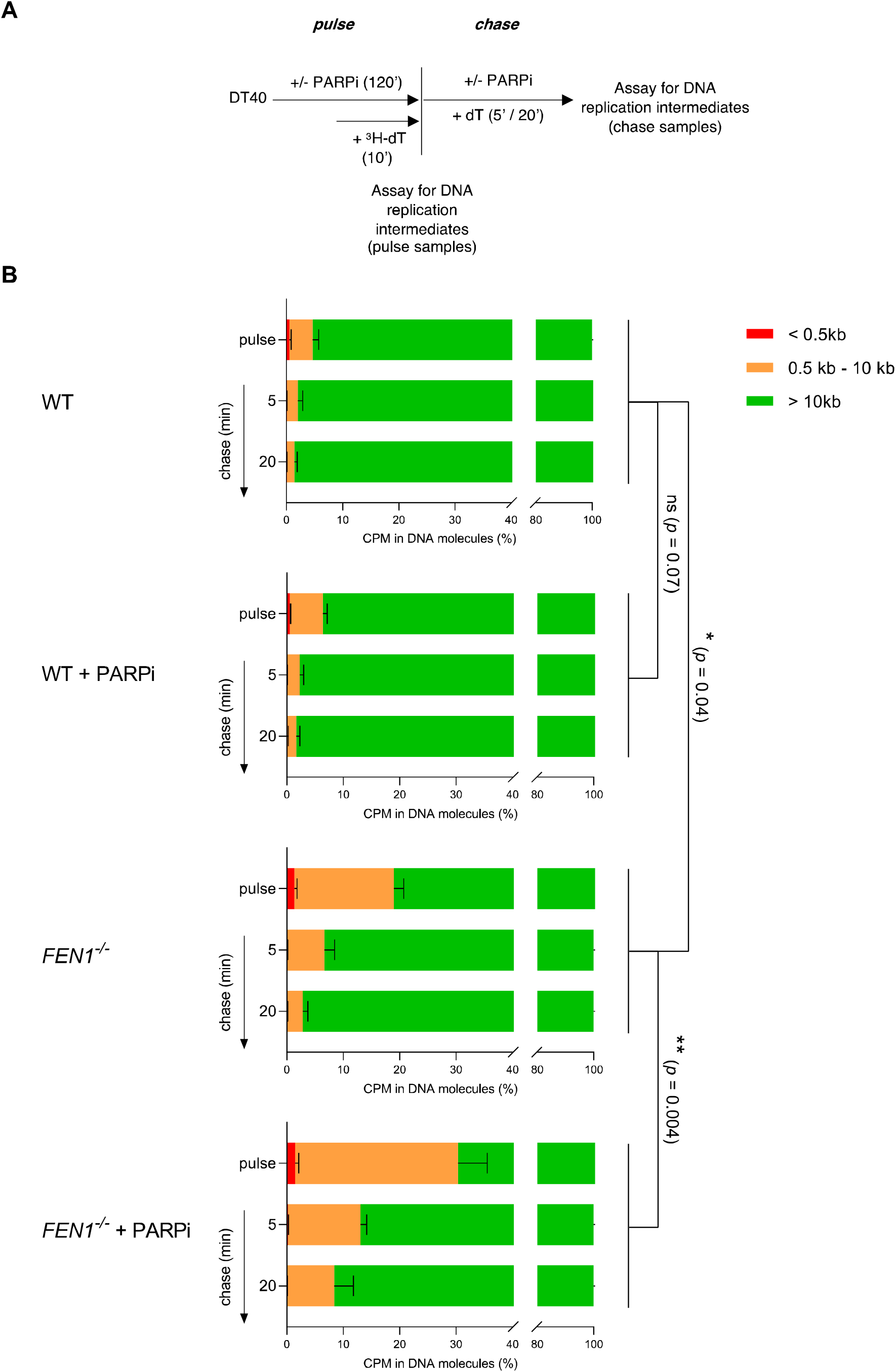
PARP inhibitor impedes the maturation of early/small (<10 kb) nascent DNA strands in *FEN1^−/−^* chicken DT40 cells. **(A)** Schematic for measuring the size distribution of ^3^H-labelled nascent DNA strands in wild type (WT) and *FEN1^−/−^* DT40 cells following 10 min pulse labelling with ^3^H-thymidine and during a subsequent 20 min cold chase, in the absence or presence of 10 μM PARPi (Olaparib) as indicated, by alkaline agarose gel electrophoresis, **(B)** Quantification of ^3^H-pulse-labelled nascent DNA strands in the size ranges <0.5 kb, 0.5-10 kb, and >10 kb by liquid scintillation counting of radioactivity in alkaline agarose gel slices. Graphs show the mean fraction (%) ±SD of ^3^H radioactivity (in CPM) in nascent DNA fragments of the indicated sizes from 3 independent experiments. Statistical analysis of the fraction of radioactivity detected as nascent DNA strands of <10 kb was conducted by 2-way ANOVA with Tukey’s post hoc multiple comparisons test.

To confirm that the impact of PARP inhibitor on nascent strand integrity reflected the induction and/or persistence of post-replicative nicks and/or gaps, rather than increased replication fork stalling and/or collapse, we employed DNA combing. DT40 cells were labelled for 15 min with CldU, followed by a further 45 min with IdU in the presence or absence of PARP inhibitor, and then subject to DNA combing to quantify the length of individual DNA replication tracts (Fig.4A). Where indicated, genomic DNA was treated with S1 nuclease prior to DNA combing, to detect post-replicative single-strand nicks and gaps. Notably, DNA replication fork rates were similar in wild type and *FEN1^−/−^* DT40 cells and were unaffected by PARP inhibitor in either cell line (Fig.4B, C & Extended Data Fig.3A, B, compare “-S1 nuclease” samples), suggesting that that PARP inhibition did not measurably affect the frequency and/or persistence of fork stalling, collapse, or reversal. However, in *FEN1^−/−^* cells, treatment with S1 nuclease reduced the median length of IdU replication tracts synthesised in the presence of PARP inhibitor by ~30%, confirming that PARP inhibition increased the number and/or persistence of post-replicative single-strand nicks/gaps if canonical Okazaki fragment processing was perturbed (Fig.4C, D & Extended Data Fig.3B, C, compare “+ S1 nuclease” samples).

**Figure 4.**
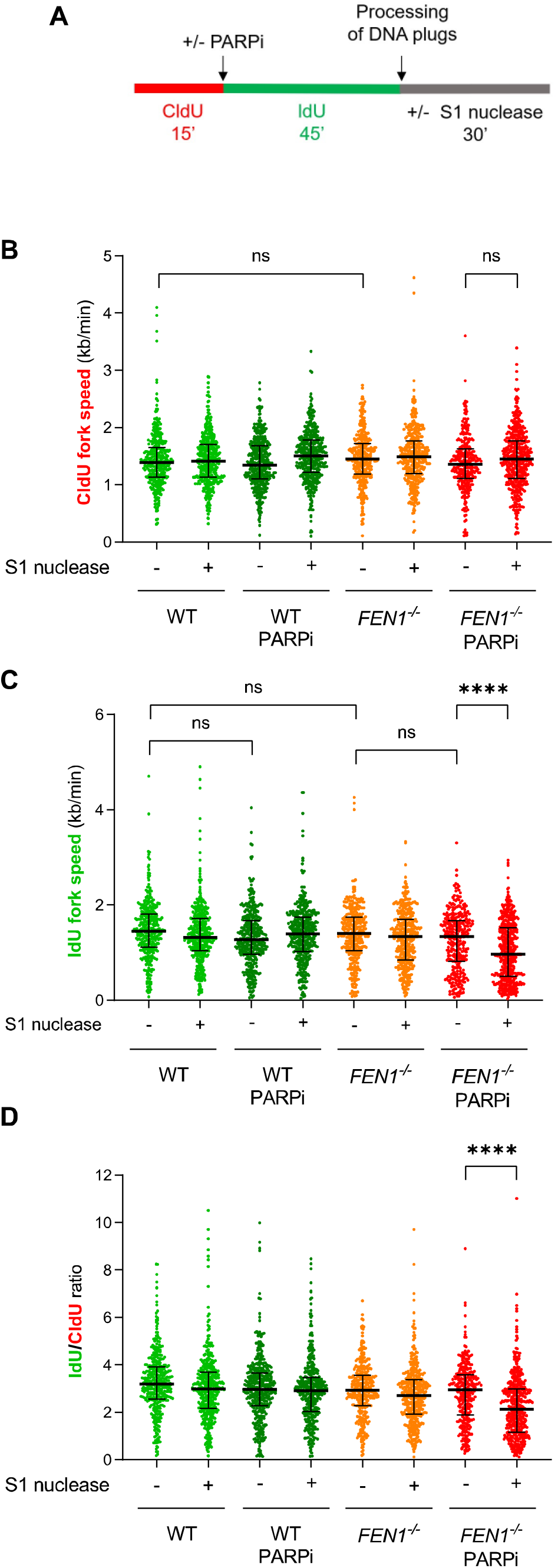
PARP inhibitor induces post-replicative single-strand gaps in *FEN1^−/−^* DT40 cells. **(A)** Schematic for measuring the rates of DNA replication fork progression during consecutive CldU and IdU pulse labelling by DNA combing in wild type (WT) and *FEN1^−/−^* DT40 cells incubated or not as indicated in 10 μM PARPi (Olaparib). **(B)** CldU tract-lengths in dual-labelled DNA fibres were measured in 3 independent experiments (>65 fibres per sample per experiment) and the distributions of CldU fork speeds (calculated assuming a constant stretching factor of 2kb/μm) are presented as scatter plots. Bars depict the median and interquartile range. Individual data points from the three experimental repeats are plotted separately in Figure S3A and statistical analysis was conducted by 2-way ANOVA (mixed effects model) with Tukey’s post hoc multiple comparisons test. **(C)** IdU fork speeds in dual-labelled DNA fibres scored and analysed as in panel B. Individual data points from the three experimental repeats are plotted separately in Figure S3B. **(D)** The ratio of IdU and CldU tract lengths in dual-labelled fibres, presented as distributions in scatter plots and analysed as in panel B. Individual data points from the three experimental repeats are plotted separately in Figure S3C. For all panels; ns, not significant; ****p<0.0001.

Collectively, our experiments with DT40 cells suggest that PARP1 is activated by unligated Okazaki fragment intermediates over large distances behind DNA replication forks, and that PARP inhibition impedes their maturation or repair. To examine whether this is also true in human cells, we disrupted FEN1 in U2OS cells by gene editing (Extended Data Fig.4). Similar to DT40 cells, *FEN1^−/−^* U2OS cells exhibited higher levels of ADP-ribosylation in S phase than did wild type cells, irrespective whether or not PARG was inhibited (Fig.5A). To examine whether this activity was located behind DNA replication forks, we compared the proximity of ADP-ribose and EdU-labelled tracts of nascent DNA immediately after pulse-labelling and following a subsequent thymidine chase, by proximity ligation assays (PLA). Whilst we detected significant PLA signal in wild type U2OS cells immediately after pulse labelling for 10 min, this signal increased significantly during a subsequent 10 min chase (Fig.5B). A similar trend was observed in *FEN1^−/−^* U2OS cells, but as expected with overall higher levels of PLA signal (Fig.5B). Importantly, the increase in PLA signal in U2OS cells during a 10 min chase did not reflect a general increase in either EdU or ADP-ribose, which was similar throughout the experiment (Fig.5C). We thus conclude that levels of S phase ADP-ribosylation are highest behind DNA replication forks, in human cells. To confirm that, similar to DT40 cells, PARP inhibitors impede the repair and/or maturation of nascent DNA strands in human cells, we employed alkaline BrdU comet assays. Indeed, similar to DT40 cells, incubation with PARP inhibitor reduced the integrity of nascent DNA strands in wild type U2OS cells during DNA replication, and did so to a greater extent in *FEN1^−/−^* U2OS cells (Fig.6A & Extended Data Fig.5A). Moreover, similar results were observed in wild type human RPE-1 cells, if we employed FEN1 inhibitor instead of FEN1 deletion (Fig.6B & Extended Data Fig.5B).

**Figure 5.**
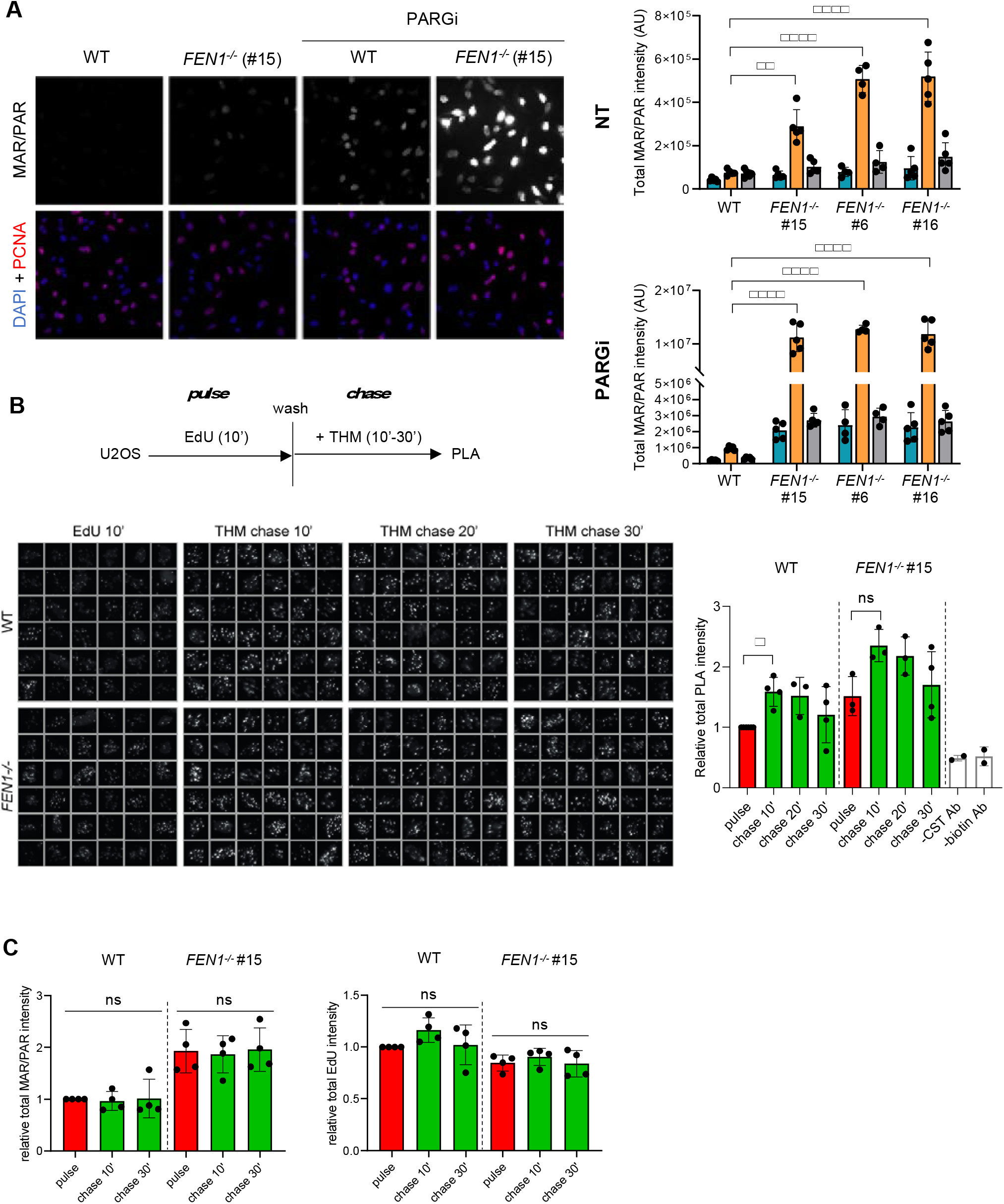
S phase PARP activity is highest behind DNA replication forks in human cells. **(A)** PARP activity in wild type (WT) and *FEN1^−/−^* U2OS cells (clones #15, #6, #16), incubated with or without PARG inhibitor (10 μM) for 30 min, measured by anti-ADP-ribose (MAR/PAR) (CST 83732) immunofluorescence in detergent extracted cells. Representative microscopy images (WT and *FEN1^−/−^* clone #15 only) are shown *left*, and ScanR quantification (WT and all *FEN1^−/−^* clones) is shown *right*. S phase cells were distinguished by positive PCNA staining and G1/G2 cells (PCNA-negative) by DNA content (measured with DAPI). Data represent the mean (±SD) total intensity of MAR/PAR (in arbitrary units (AU)) from 5 independent experiments with individual data points plotted. Statistical significance was assessed by 1-way ANOVA with post-hoc Sidak’s multiple comparisons test. **(B)** Physical proximity of newly incorporated EdU and ADP-ribose (MAR/PAR; CST 83732) in wild type (WT) and *FEN1^−/−^* U2OS cells, measured following pulse labelling for 10 min and during a subsequent 30 min thymidine (THM) chase by PLA, as indicated in the schematic. Sampled cells were pre-extracted and fixed using PFA. PLA was measured using anti-biotin antibodies to detect biotin-azide clicked to EdU and anti-ADP-ribose antibodies to detect sites of PARP activity. Representative ScanR image galleries (*left*, each box is a single cell) and quantification (*right*) are shown. Total PLA intensity in PCNA positive cells was normalized to that in wild type U2OS cells immediately after EdU pulse. Data represent mean (±SD) from 3-4 independent experiments with individual data points plotted. Statistical significance was assessed by 1-way ANOVA with post-hoc Sidak’s multiple comparisons test. **(C)** Quantification of ADP-ribose and EdU levels following pulse labelling and during the thymidine chase. Cells were treated and fixed as in panel A and EdU and ADP-ribose intensities were determined by ScanR in PCNA positive cells. Data represent mean (±SD) from 3-4 independent experiments with individual data points plotted. Statistical significance was assessed by 1-way ANOVA with post-hoc Sidak’s multiple comparisons test. For all panels; ns, not significant, *p< 0.05.

**Figure 6.**
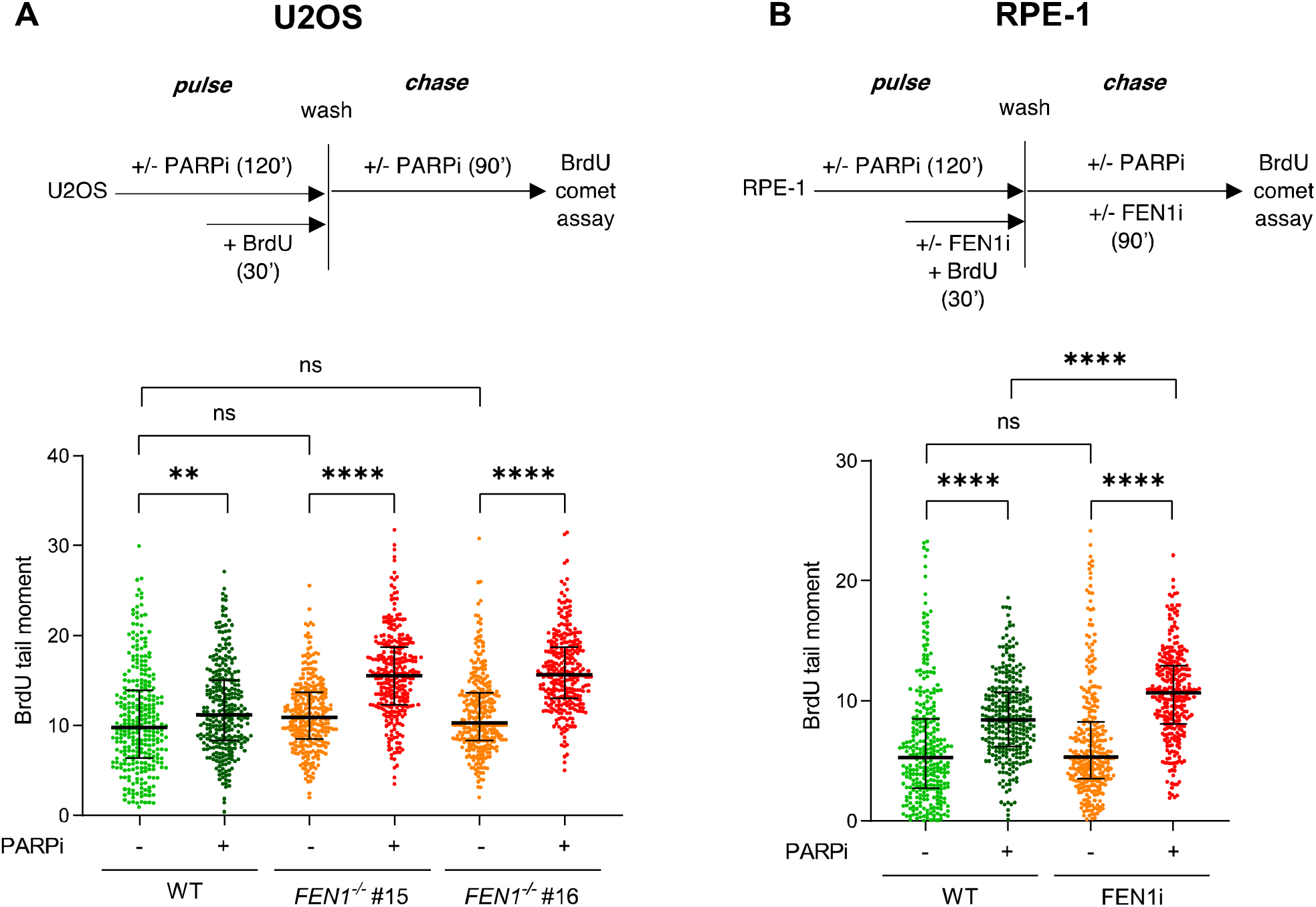
PARP inhibitor impedes the maturation of nascent DNA strands in human cells. (A) DNA breaks in nascent DNA strands quantified in wild type (WT) and *FEN1^−/−^* (clones #15 & #16) U2OS cells by alkaline comet assays after pulse labelling (30 min) with BrdU followed by a subsequent chase (90 min) as indicated in the schematic (*top*). 10 μM PARPi (Olaparib) was employed or not as indicated. Nascent DNA strands were scored for comet tail moments by staining with anti-BrdU antibodies. For each sample, scatter plots are anti-BrdU comet tail moments from 300 cells combined from 3 independent experiments (100 cells per sample per experiment) and bars are the median and interquartile range. The individual data sets from the three experimental repeats are plotted separately in Figure S5A, and statistical analysis was conducted as in Figure 2A. **(B)** DNA breaks in nascent DNA strands quantified in wild type RPE-1 cells by alkaline comet assays after pulse labelling (30 min) with BrdU followed by a subsequent chase (90 min) as indicated in the schematic (*top*). Where indicated, cells were incubated with 10 μM PARPi (Olaparib) and/or 10 μM FEN1 inhibitor (FEN1i). Nascent DNA strands were scored for comet tail moments by staining with anti-BrdU antibodies. Scatter plots are as in panel A, with separate experimental repeats plotted in Figure S5B. For all panels; ns, not significant; **p<0.01, ****p<0.0001.

In summary, we show here that PARP activity is greatest behind DNA replication forks, and that PARP inhibitors impede the maturation of nascent DNA strands during DNA replication. Moreover, the impact of PARP inhibition on nascent strand integrity is particularly pronounced in cells lacking FEN1, implicating unligated Okazaki fragments as an endogenous source of PARP inhibitor-induced genotoxicity.

## Discussion

PARP inhibitors provide a powerful new approach in the treatment of cancer, particularly in tumour cells in which HR is attenuated or absent ^32–34^. By trapping PARP1 on DNA lesions PARP inhibitors impede DNA repair and render proliferating cells dependent on HR for cell survival. However, the identity of the DNA structures on which PARP1 becomes trapped and that exert this cytotoxicity have been unclear ^12^. Here, we have found that PARP inhibitors slow the maturation of nascent DNA strands during DNA replication. It is unlikely that this finding is explained by an impact of PARP inhibitor on DNA replication fork progression, resulting from the role identified for PARP1 in regulating replication fork reversal and/or repair following treatment of cells with genotoxins ^20,21,36^. This is because PARP inhibitor did not affect DNA replication fork rates in our experiments, as measured by DNA combing, suggesting that the impact of PARP1 in regulating replication fork progression was too rare to be detected in unperturbed cells. This result contrasts with a recent report in which PARP inhibition increased replication forks speeds in unperturbed cells ^37^, perhaps reflecting the use of longer periods of PARP inhibition in the latter study.

In contrast to the lack of impact on DNA replication fork rates, our DNA combing experiments did detect an increased number of post-replicative single-strand gaps in *FEN1^−/−^* cells following treatment with PARP inhibitor. This is in agreement with the greatly exacerbated impact of PARP inhibitor on nascent strand maturation detected in these mutant cells by alkaline comet assays and alkaline gel electrophoresis. Although FEN1 has multiple roles in DNA metabolism the most common role of this nuclease is processing Okazaki fragment intermediates ^38^, which is consistent with the S phase-specific impact of PARP inhibitor in *FEN1^−/−^* cells in our experiments. *FEN1^−/−^* DT40 cells are hypersensitive to PARP inhibitor ^35^, and in our clonogenic experiments these cells were far more sensitive to PARP inhibitor than were HR-deficient cells lacking XRCC3. Collectively, our data argue that unligated Okazaki fragments are a major source of PARP1 trapping and nascent strand breakage, in the presence of PARP inhibitor.

Our alkaline comet experiments suggest that PARP1 becomes trapped primarily on later/large intermediates of Okazaki fragment maturation, unless canonical Okazaki fragment processing is also perturbed following which early/small maturation intermediates are also affected. This is because in wild type cells we detected an impact of PARP inhibitor in alkaline comet assays, which in our hands are sensitive to DNA fragments of >500 kb, but not by gel electrophoresis assays that can detect fragments of <10kb. In contrast, in *FEN1^−/−^* DT40 cells, PARP1 inhibitor reduced the integrity of nascent DNA strand in both assays. Thus, our findings suggest that in wild type cells PARP inhibitors primarily slow the maturation of nascent DNA strands tens-to-hundreds of kilobases behind DNA replication forks, and do so additionally much closer to DNA replication forks if canonical Okazaki fragment processing is also perturbed. This idea is consistent with our previous observations in RPE-1 cells using confocal microscopy, in which focal sites of PCNA and S phase ADP-ribosylation were typically adjacent to each other rather than overlapping, unless FEN1 was inhibited ^13^. The idea that PARP activity is normally greatest at considerable distances behind DNA replication forks is also consistent with our PLA data, which revealed that sites of EdU-labelled nascent DNA were nearer to sites of ADP-ribosylation following a 10 min chase than immediately after 10 min pulse labelling. Interestingly, the PLA signal appeared to decline thereafter in our experiments, perhaps reflecting the loss of ADP-ribosylation at sites of pulse labelled nascent DNA as the latter are repaired, although this decline did not reach statistical significance.

The cytotoxicity of PARP inhibitors reflects, in part at least, the trapping of PARP1 on DNA breaks, which impedes their repair by other DNA repair enzymes ^34,35^. Our data argue strongly that unligated intermediates of Okazaki fragment processing comprise a large fraction of the DNA structures on which PARP1 is trapped by PARP inhibitors, in S phase. Whether or not PARP1 plays an active role in processing Okazaki fragment intermediates, or whether it simply becomes trapped on these structures in the presence of PARP inhibitor, remains to be determined. However, our current and previous observations that the level of PARP1 activity is highest in S phase and is increased further if canonical Okazaki fragment processing is perturbed by FEN1 inhibition, FEN1 deletion, or LIG1 mutation is consistent with the former, as is the observation that SSB repair proteins such as XRCC1 are recruited to sites of PARP1 activity in S phase ^13,39–41^. Indeed, it has been reported that the XRCC1 partner protein LIG3 can replace LIG1 during Okazaki fragment processing in DT40 cells, and in *Xenopus* extracts ^29,30,42^. However, irrespective of whether PARP1 is an active participant or bystander during Okazaki fragment processing, our data identify these and possibly other nascent strand intermediates as major sources of genome breakage and cytotoxicity in cells following treatment with PARP inhibitors. Our data thus shed light on the source of DNA breaks that might underpin the clinical utility of PARP inhibitors in cancer therapy.

## Methods

### Chemicals and antibodies

100 mM stock solution of BrdU (Merck, B5002) and 10 mM stock solutions of PARG inhibitor (PDD 0017273 - Tocris, 5952; Merck, SML1781), PARP inhibitors (KU 0058948 - Axon Medchem, Axon 2001; Olaparib - ApexBio, A4154), FEN1 inhibitor (synthesized as described in Tumey et al., 2005, compound 17), and EdU (Cambridge Bioscience, CAY20518) were prepared in dimethyl sulfoxide (DMSO). CldU (Merck, C6891) and IdU (Merck, I7125) were dissolved directly in culture medium at a final concentration of 2.5 mM, and thymidine (Merck, T1895) in culture medium at 200 mM. 1 mCi/ml ^3^H-Thymidine (PerkinElmer, NET027W005MC) in 2% ethanol was added directly to culture medium to a final concentration of 2 μCi/ml.

Primary antibodies used are as follows: Anti-poly-ADP-ribose binding reagent/ PAR reagent (recombinant protein fused to rabbit Fc tag; Millipore, MABE1031), Rabbit monoclonal anti-Poly/Mono-ADP Ribose (anti-PAR/MAR, Cell Signaling, 83732), Mouse monoclonal anti-PCNA (Santa Cruz, sc-56), Rabbit monoclonal anti-PARP1 (Cell Signaling, 9532), Rat polyclonal anti-α-tubulin (Abcam, ab6160), Rabbit polyclonal anti-H3 (Abcam, ab1791), Rabbit polyclonal anti-FEN1 (LifeSpan Biosciences, LS-C80825), Mouse monoclonal anti-biotin (Merck, BN-34), Mouse monoclonal anti-PARP1 (Santa Cruz, sc-8007), Rat recombinant anti-PCNA (Abcam, ab252848), Rat monoclonal anti-BrdU (Abcam, ab6326), Mouse monoclonal anti-BrdU (Becton Dickinson, 347580), Mouse monoclonal anti-ssDNA (Millipore, MAB3034).

Secondary antibodies used are as follows: HRP-conjugated goat anti-rabbit (Bio-Rad, 170-6515), HRP-conjugated goat anti-mouse (Bio-Rad, 170-6516, HRP-conjugated rabbit anti-rat (Abcam, ab6734), Donkey anti-rabbit Alexa Fluor 488 (Thermo Fisher, A21206), Donkey anti-mouse Alexa Fluor 568 (Thermo Fisher, A10037), Donkey anti-mouse Alexa Fluor 647 (Thermo Fisher, A31571), Goat anti-mouse Alexa Fluor 488 (Thermo Fisher, A11001), Goat anti-mouse Alexa Fluor 488 (Thermo Fisher, A32723), Donkey anti-goat Alexa Fluor 488 (Thermo Fisher, A11055), Goat anti-rat Alexa Fluor 568 (Thermo Fisher, A11077), Donkey anti-rat Alexa Fluor 488 (Thermo Fisher, A21208).

### Cell culture

Human hTERT RPE-1 cells (ATCC, CRL-4000) were maintained in Dulbecco’s Modified Eagle’s Medium (DMEM/F12, Merck) supplemented with 10% fetal calf serum (FCS). Human wild type (ATCC, HTB-96) and *FEN1^−/−^* U2OS cells were cultured in DMEM (Gibco) with 10% FCS and 2 mM L-glutamine (Gibco). Wild type and *FEN1^−/−^* U2OS cells were grown under 3 % oxygen levels. Chicken wild type, *FEN1^−/−^* ^31^ and *XRCC3^−/−^* ^46^. DT40 cell lines (a gift from S. Takeda) were cultured in RPMI 1640 medium supplemented with 10% FCS, 1% chicken serum (Gibco), 2 mM L-glutamine, 10 μM β-mercaptoethanol (Gibco). All growth media was supplemented with penicillin (100 Units/ml)/streptomycin (100 mg/ml) (Merck) and all cells were grown at 37°C. U2OS and *FEN1^−/−^* cells were grown under 3 % oxygen levels.

### Purification of SpCas9 and generation of *FEN1^−/−^* U2OS cells

His-SpCas9-GFP was expressed in and purified from BL21 (DE3, NEB, C2527H) bacteria as previously described ^47^. Briefly, inoculated culture was grown to OD_600_ 0.5, cooled down to 16°C and induced by 0.1 mM IPTG and incubated for 20 h. Cells were resuspended in lysis buffer (50 mM Tris pH 7.5, 500 mM NaCl, 20 mM imidazole, 1 mM TCEP) supplemented with protease inhibitors, sonicated and centrifuged at 20,000 g for 40 min at 4°C. Supernatant was incubated with Ni-NTA agarose beads (GE Healthcare, 17-5318-01) for 1 h at 4°C, beads were extensively washed with lysis buffer and then lysis buffer containing 150 mM NaCl. His-SpCas9-GFP was eluted with 300 mM imidazole in 50 mM Tris pH 7.5, 150 mM NaCl, 1 mM TCEP, diluted with 25 mM HEPES pH 7.4, 150 mM NaCl, 1 mM TCEP and loaded to 5 ml HiTrap SP HP (GE Healthcare, 17-1152-01). After extensive washing, elution was done using gradient to 60% in 25CV followed by 8CV to 100% (1 M NaCl), 2.5 ml fractions were captured and snap frozen in liquid nitrogen before use. To generate Cas9 RNPs for electroporation, 120 pmol crRNA (Merck, UGUGGCCCCCAGUGCCAUCC) was mixed with 120 pmol tracrRNA (Merck, TRACRRNA05N) in 1:1 molar ratio in Cas9 buffer (20 mM HEPES pH 7.5, 150 mM NaCl, 2 mM MgCl2, 1 mM TCEP before addition of 100 pmol His-Cas9-GFP and incubation for 10 min at room temperature. 2×10^5^ U2OS cells were washed in PBS and electroporated using Neon transfection system (Thermo Fisher) with 10 μl tip using 1230 V / 10 width / 4 pulses settings. After 3 days cells were reseeded to 96 well plate at 0.5 cell/well. Single cell clones were analyzed by Western blotting, genomic DNA was isolated (DNeasy Blood & Tissue Kit, Qiagen, 69504), locus surrounding Cas9 cutting site was amplified using Q5 DNA polymerase (NEB, M0491S) and primers around the Cas9 cut site (FWD: TGGTGCCGCGCGGCAGCCACCTGTCTTTCAGGTCTGCCAT, REV: CACCAGTCATGCTAGCCATATTCACTGGCAGTCAGGTGTC), PCR products were purified (QIAquick PCR Purification Kit, Qiagen, 28106), cloned into NdeI cut pET28a using NEBuilder HiFi DNA Assembly Master Mix (NEB, E2621S), single colonies picked and Sanger sequenced.

### PLA

Cells were seeded at 2×10^5^ cells per well of 6-well plate. Next day, cells were incubated with 100 μM EdU (Cambridge Bioscience, CAY20518) for 10 min followed by 3 washes and incubation in media containing 100 μM thymidine (Merck). Before fixation, cells were washed with PBS, pre-extracted using pre-extraction buffer (25 mM HEPES pH 7.4, 50 mM NaCl, 1 mM EDTA, 3 mM MgCl_2_, 0.3 M sucrose, 0.5% Triton X-100) supplemented with 10 μM PARPi (KU0058948, Axon Medchem, Axon 2001) and PARGi (PDD0017273, Merck, SML1781) 5 min on ice and fixed with cold 4% formaldehyde for 15 min. Cells were permeabilized using ice cold methanol/acetone solution (1:1) for 5 min and PBS containing 0.5% Triton X-100 and blocked in BSA. Click reaction was performed using 0.1 M Tris pH 8.5, 0.1 M sodium ascorbate, 2 mM Cu_2_SO_4_ and 0.1mM biotin-azide (Merck, 762024) or AlexaFluor 647 azide (Thermo Fisher, A10277) for 45 min at room temperature. Cells were washed and stained with indicated primary antibodies for 2 h at RT followed by incubation with PLA probes (Merck, Duolink® In Situ PLA® Probe Anti-Rabbit PLUS, DUO92002, Duolink® In Situ PLA® Probe Anti-Mouse MINUS, DUO92004) for 1 h at 37 °C, ligation for 30 min 37 °C and polymerase reaction overnight at 37 °C according to manufacturer’s protocol (Merck, Duolink® In Situ Detection Reagents Red, DUO92008). Images were acquired using an Olympus IX81 microscope equipped with ScanR Screening System using 40x objective at a single autofocus-directed z-position under non-saturating settings. The inbuilt Olympus ScanR Image Analysis Software was used to analyse acquired images. Nuclei were identified by DAPI signal using an integrated intensity-based object detection module. Fluorescence intensities of interest were quantified in PCNA positive cells.

### Detection of ADP-ribose levels by indirect immunofluorescence

U2OS cells were seeded at 2×10^5^ cells per well of 6-well plate. Next day, cells were treated with 10 μM PARG inhibitor (PDD0017273, Merck, SML1781) for 30 min. Before fixation, cells were washed with PBS, pre-extracted using pre-extraction buffer (25 mM HEPES pH 7.4, 50 mM NaCl, 1 mM EDTA, 3 mM MgCl_2_, 0.3 M sucrose, 0.5% Triton X-100) supplemented with 10 μM PARPi (KU0058948, Axon Medchem, Axon 2001) and PARGi (PDD0017273, Merck, SML1781) 5 min on ice and fixed with cold 4% formaldehyde for 15 min. Cells were permeabilized using ice cold methanol/acetone solution (1:1) for 5 min and PBS containing 0.5% Triton X-100 and blocked in BSA. Cells were stained with indicated primary antibodies for 2 h at RT followed by incubation with secondary antibodies for 1 h at RT, after washing in PBS, DNA was stained using DAPI. DT40 cells were collected, washed, and diluted in ice-cold PBS to a final concentration of ~7×10^5^ cells/ml. The cell suspension was centrifuged on a microscope slide (Thermo Fisher Scientific) (200 μl/slide) at 800 rpm for 3 min in a Cytospin centrifuge, and PAP Pen Liquid Blocker (Merck) was used to draw a circle around a specimen to hold reagents within the area containing cells. Then, cells were fixed with 4% formaldehyde in PBS for 10 min at RT, rinsed in PBS and subsequently permeabilized with ice-cold methanol/acetone solution (1:1) for 5 min at RT, followed by 3 short washes in PBS. Next, cells were incubated in the blocking solution (3% BSA in PBS) for 1 h at RT, followed by incubation with appropriate primary antibodies (1 h at RT) and then with fluorochrome-conjugated secondary antibodies (1 h at RT). Slides were washed 3 times in PBS after both primary and secondary antibody incubations. Next, DNA was stained with DAPI (1 mg/ml in water) for 5 min at RT, slides were washed in water, and cells were mounted in fluoroshield (Merck). Immunofluorescence images were acquired using an Olympus IX81 microscope equipped with scanR Screening System using 40x objective at a single autofocus-directed z-position under non-saturating settings. The inbuilt Olympus scanR Image Analysis Software was used to analyse acquired images. Nuclei were identified by DAPI signal using an integrated intensity-based object detection module. The G1, S and G2 phase cells were gated based on PCNA and DAPI intensity and fluorescence intensities of interest were quantified. High-resolution images in Figure 1A were acquired with an Apotome widefield microscope (Zeiss) using 63x oil objective.

### Chromatin fractionation assay

DT40 cells (~5×10^6^/sample) were harvested and lysed for 20 min on ice in 200 μl of CSK buffer (25 mM HEPES pH 7.4, 150 mM NaCl, 0.3 M sucrose, 3 mM MgCl2, 1 mM EDTA, 0.5% Triton X-100) containing protease inhibitors (Roche) and phosphatase inhibitors (Merck), 50 μl of samples were collected (total cell lysates). Soluble and chromatin-bound proteins were separated by centrifugation (5 min 20,000 g at 4 °C) and supernatants were collected (soluble fractions). Pellets were washed twice in 1 ml of CSK buffer and were dissolved in 150 μl of 2x Laemmli sample buffer (chromatin fractions). The following steps were the same as for western blotting (see below).

### Western blotting

Cells were lysed in 2x Laemmli buffer lacking reducing agent (100 mM Tris-HCl pH 6.8, 4% SDS, 20% glycerol) followed by incubation at 99°C for 5 min and sonication. Protein was quantified using BCA assays (Thermo Fisher Scientific), DTT and bromophenol blue added to 0.1M and 0.1% respectively, and samples heated for 10 min at 99 °C. Samples were resolved on Bis-Tris SDS-PAGE gels in MOPS buffer (pH 7.7, 100-150 V) and transferred to nitrocellulose membrane (Thermo Fisher Scientific). Membranes were blocked for 1 h in 1x TBS containing 0.1% Tween20 (TBST) and 5% milk, followed by incubation with appropriate primary antibodies either for 1 h at RT or overnight at 4 °C. Then, membranes were incubated with the horseradish peroxidase-conjugated secondary antibodies for 1 h at RT. After both primary and secondary antibody incubations membranes were washed 3×10 min in TBST at RT. ECL detection reagent (GE Healthcare or Thermo Fisher Scientific) was applied and immunoreactive proteins were visualized either using ImageQuant LAS 4000 machine (Raytek) or chemiluminescence film (Scientific Laboratory Supplies or GE Healthcare).

### Clonogenic survival assay

WT, *FEN1^−/−^* and *XRCC3^−/−^* DT40 cells were seeded in triplicate in 6-well plates at 100, 500, or 2500 cells/well depending on PARPi dose in 5 ml of medium supplemented with 1.5% methylcellulose (Merck) and the indicated concentrations of Olaparib. Cells were grown for 10-14 days at 37°C and visible colonies counted. Survival (%) was defined as the average number of colonies on treated plates divided by the average number of colonies on untreated plates multiplied by 100.

### Alkaline comet assays

Alkaline comet assays were performed essentially as described (Breslin et al., 2006). For measuring DNA breaks in total genomic DNA, slides were stained with SYBR Green (Merck, 1:10,000) or with propidium iodide (PI - Merck, 1:500), and with p-Phenylenediamine dihydrochloride (Thermo Fisher Scientific, 41 μg/ml) in PBS as an antifade. To detect S phase cells and to detect DNA breaks specifically in nascent strands cells were pulse-labelled with 100 μM BrdU for 30-45 min as indicated, and then ether sampled immediately (to detect S phase cells) or incubated for a subsequent 90 min chase period (to measure breaks in nascent strands during the maturation of DNA replication intermediates). After the neutralisation step, slides were washed 3×10 min in PBS, followed by incubation with the mouse monoclonal anti-BrdU antibody (Becton Dickinson, 347580; 1:2) overnight at 4 °C in a humid chamber. Excess primary antibody was removed, slides were then incubated simultaneously with two different secondary antibodies diluted in PBS/0.1% Tween 20/3% BSA for 1 h at RT to amplify the signal (goat anti-mouse Alexa Fluor 488 (Thermo Fisher, A11001; 1:250) and donkey anti-goat Alexa Fluor 488 (Thermo Fisher, A11055; 1:250)). Thereafter, slides were washed 3×10 min in PBS and counterstained with PI (Merck, 1:500) and p-Phenylenediamine dihydrochloride (Thermo Fisher Scientific, 41 μg/ml) in PBS. In all cases, the lysis buffer was pH 10.4. Comet tail moments were visualised using Nikon Eclipse 50i widefield microscope and scored with Comet Assay IV software (Perceptive Instruments) in SYBR Green (with GFP filter) or PI (with FITC filter) labelled DNA for total genomic DNA breaks, and comet tail moments in anti-BrdU-stained DNA (with GFP filter) were scored for DNA breaks in DNA nascent strands.

### Alkaline agarose gel electrophoresis

Analysis of nascent DNA fragments by agarose gel electrophoresis was conducted as described ^29^. DT40 cells (~5×10^6^/sample) were pulse-labelled with ^3^H-thymidine (2 μCi/ml) for 10 min, followed by 5-20 min chase in fresh medium containing 2 mM thymidine. Cells were collected, washed in ice-cold PBS, and resuspended in 20 μl of Buffer A (10 mM Tris–HCl, pH 8.0; 50 mM NaCl; 0.1 M EDTA). Next, the cell suspension, prewarmed for 10 sec at 50 °C, was gently mixed with 25 μl of molten 1.5 % low-melting-point agarose and pipetted into a casting mold (Bio-rad), which was placed on ice for 5 min to let agarose plugs solidify. Subsequently, plugs were lysed in 1ml of Buffer A containing 0.2 mg/ml proteinase K (Thermo Fisher Scientific) and 2% N-lauryl sarcosine (Merck) for 18 h at 50°C, followed by washing in 5 ml of Buffer A for 1 h at RT. The agarose plugs were then loaded on the comb, embedded in 1% alkaline agarose gel (1% agarose, 50 mM NaOH, 1 mM EDTA in H_2_O) and the genomic DNA fractionated by electrophoresis under denaturing conditions (50 mM NaOH, 1 mM EDTA in H_2_O) for 7.5 h (2 V/cm) at RT. Following electrophoresis, the gel was neutralised for 1 h at RT in 1M Tris–HCl, pH 7.6/1.5 M NaCl and stained with SYBR Green (Merck) (1:10,000) to visualise DNA molecular mass markers (0.075 to 20 kb). For each sample lane, the gel was cut into 1 cm length slices that were placed in scintillation vials and soaked in 0.1 M HCl for 1 h. The HCl solution was then carefully removed and the gel slices melted in a microwave. 4 ml of aqueous scintillant was thoroughly mixed with the melted gel slices by vortexing and ^3^H quantified (counts per minute; CPM) in a scintillation counter. The radioactivity in agarose slices corresponding to fragment sizes of <0.5 kb, 0.5 −10 kb, and >10 kb were combined and plotted as percentages of the total CPM in all gel slices of that sample.

### DNA combing

DT40 cells were labelled with 25 μM CldU for 15 min, followed by labelling with 250 μM ldU for another 45 min in the presence or absence of the PARP inhibitor Olaparib (10 μM). Next, cells were washed 2 times and resuspended in ice-cold PBS at ~5×10^6^ cells/ml. 50 μl of cell suspension, prewarmed for 10 sec at 50 °C, was gently mixed with an equal volume of molten 1.5 % low-melting-point agarose and pipetted into a casting mold (Bio-rad), which was placed on ice for 10 min to solidify the agarose. The agarose plugs were then incubated in round-bottom 10 ml tubes containing 0.5 ml proteinase K solution (2 mg/ml proteinase K, 10 mM Tris-HCl pH 7.5, 100 mM EDTA, 0.5% SDS, 20 mM NaCl) overnight at 50°C, and then washed twice for 1 h each in TE50 solution (10 mM Tris-HCl pH 7.5, 50 mM EDTA, 0.5% SDS, 100 mM NaCl), 2X 1 h in TE solution (10 mM Tris-HCl pH 7.5, 1 mM EDTA, 100 mM NaCl), and then incubated in 1 ml of MES solution (35 mM MES hydrate, 150 mM MES sodium salt, 100 mM NaCl) for 20 min at 68 °C. The tubes were cooled at 42 °C for 10 min before addition of 3 μl of β-agarase (NEB) dissolved in 100 μl MES solution, and were incubated overnight at 42 °C. The agarase-treated samples were then carefully poured into combing reservoirs containing 1.2 ml of MES solution supplemented with 2 mM Zn(O_2_CCH_3_)_2_ and either S1 nuclease (40 U/ml) or S1 nuclease dilution buffer (Thermo Fisher) and were incubated for 30 min at RT. Next, the genomic DNA was combed onto silanized coverslips (Genomic vision) using a combing machine (Genomic vision), and coverslips were baked for 2 h at 60 °C. DNA was denatured in fresh 0.5 M NaOH solution containing 1 M NaCl for 8 min at RT, and coverslips were washed 3 times for 3 min in PBS. After that, coverslips were incubated in the blocking solution (1% BSA with 0.1% Tween20 in PBS) for 30 min at RT and were subsequently stained with antibodies at 37 °C in a humid chamber. First, coverslips were incubated with primary rat monoclonal anti-BrdU (Abcam, ab6326; 1:50) and mouse monoclonal anti-BrdU (Becton Dickinson, 347580; 1:25) antibodies for 1 h, followed by incubation with secondary goat anti-mouse Alexa Fluor 488 (Thermo Fisher, A11001; 1:25) and goat anti-rat Alexa Fluor 568 (Thermo Fisher, A11077; 1:25) antibodies for 45 min. Then, coverslips were incubated with mouse monoclonal anti-ssDNA antibody (Millipore, MAB3034; 1:25) for 2 h to stain all genomic DNA and subsequently with donkey anti-mouse Alexa Fluor 647 (Thermo Fisher, A31571; 1:25)) for 45 min. Coverslips were washed 3 times for 3 min in PBST after each antibody incubation. Finally, coverslips were dried and then mounted onto microscope slides in fluoroshield (Merck). High-resolution images were acquired with an Apotome widefield microscope (Zeiss) using either 40x or 63x oil objectives. ImageJ64 software (NIH, https://imagej.nih.gov/ij/) was used to measure lengths of labelled replication tracks. The speed of replication fork progression was calculated assuming a constant stretching factor of 2 kb/μm.

### Flow cytometry

DT40 cells (~2×10^6^/sample) were collected, washed, and resuspended in 100 μl of ice-cold PBS. Next, 900 μl 70% ethanol was added to the cell suspension dropwise while gently vortexing, and samples were incubated in fixing solution overnight or longer at 4 °C. Before analysis, cells were washed in PBS and stained in the dark with 500 μl of PBS solution (2 mM MgCl2, 50 μg/ml PI, 50 μg/ml RNase A) for 20 min at 37 °C. Cells were counted using BD Accuri C6 Plus Flow Cytometer. The data were analysed and visualised using FlowJo software (FlowJo LLC, https://www.flowjo.com/).

### Statistical analysis and graphs

All statistical analysis and visualisation of data were carried out in GraphPad Prism (version 9.1, https://www.graphpad.com/) unless stated otherwise. When statistical comparison of two groups was required, a two-tailed paired t-test was applied. In the cases where statistical comparison of more than two groups needed to be performed, a 1-way or 2-way Analysis of Variance (ANOVA) test was done, depending on whether the effect of one or two different factors was considered, respectively. When statistically significant differences were detected with the ANOVA test, post-hoc multiple comparisons test recommended by the software was performed. Where possible, hierarchical/nested analyses were conducted with experimental repeats as matched/blocked factors and individual comets/DNA fibre lengths as technical repeats.

**Extended Data Figure 1.**
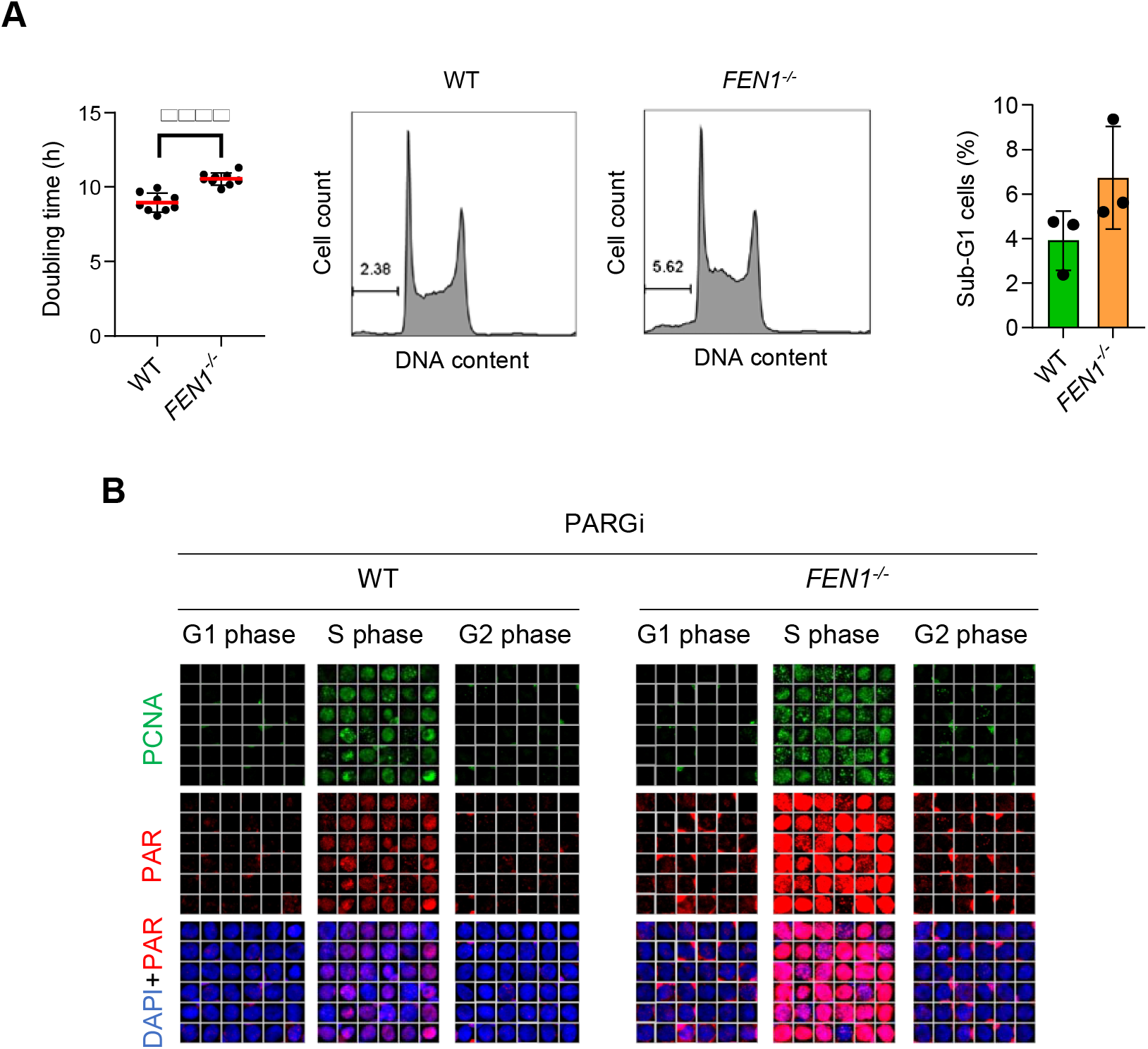
Increased doubling time and S phase poly(ADP-ribose) levels in *FEN1^−/−^* DT40 cells. **(A)** *Left*, individual data points and mean (±SD) doubling time for wild type (WT) and *FEN1^−/−^* DT40 cells from 9 independent experiments. Statistical significance was assessed by paired t test. *Middle*, representative cell cycle distribution profiles. *Right*, quantification of the fraction of cells with sub-G1 content. Data represent the mean (±SD) from 3 independent experiments with individual data points plotted. **(B)** Representative ScanR image galleries of data quantified in Figure 1A, of anti-poly(ADP-ribose) immunofluorescence (PAR) (MABE1031) in wild type (WT) and *FEN1^−/−^* DT40 cells (only PARGi-treated cells are shown).

**Extended Data Figure 2.**
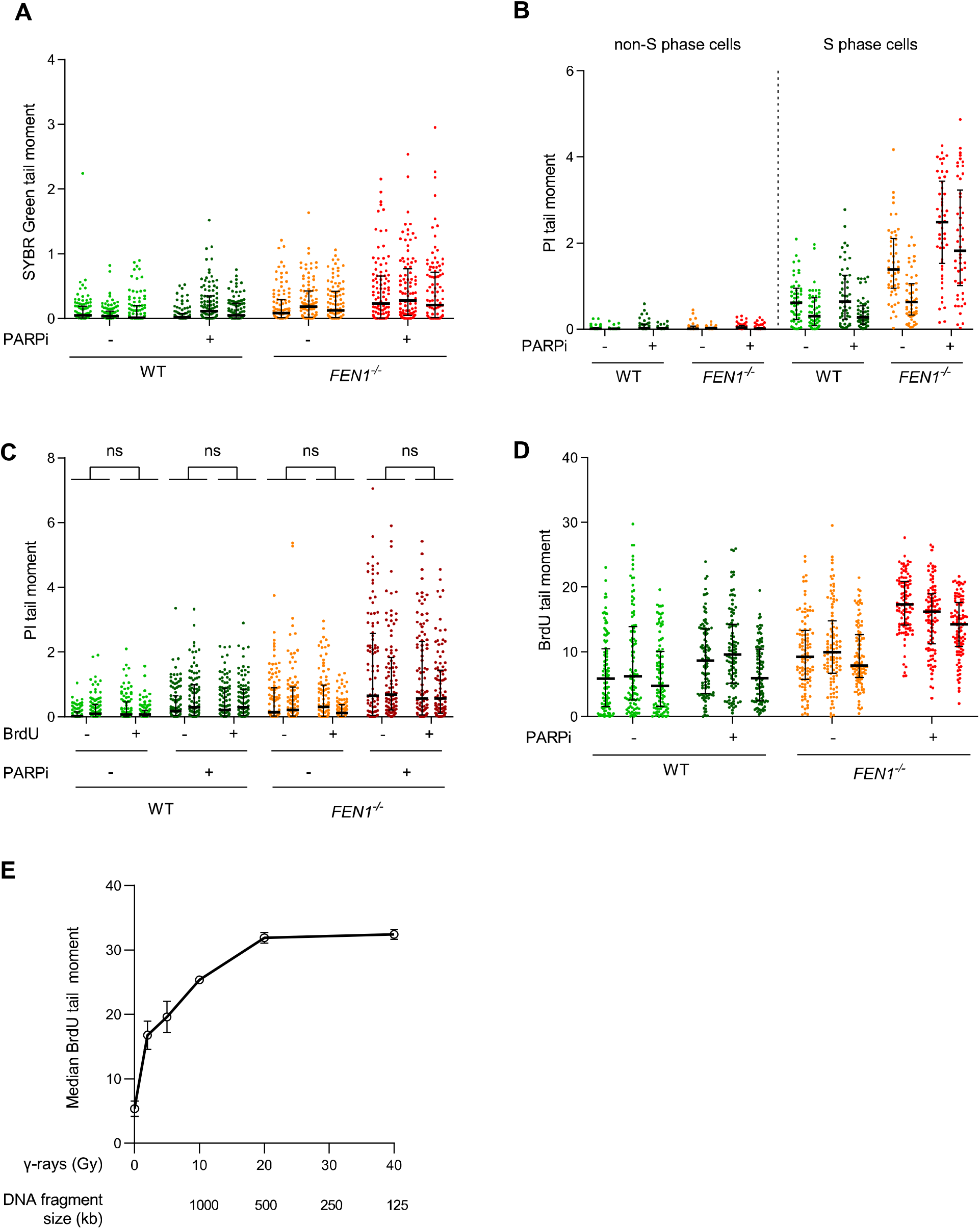
PARP inhibitor impedes the maturation of large/late nascent DNA strands in wild type and *FEN1^−/−^* chicken DT40 cells. **(A, B)** Scatter plots of the data from Figure 2A & 2B, respectively, with each experimental repeat plotted side by side. **(C)** DNA strand breaks quantified by alkaline comet assays in wild type (WT) and *FEN1^−/−^* DT40 cells pulse-labelled or not with BrdU for the last 45 min of a 2h-incubation with DMSO vehicle or 10 μM PARPi (Olaparib). Genomic DNA was visualised for scoring comet tail moments by staining with propidium iodide (PI). Plotted data are the individual comet tail moments from 2 independent experiments (100 cells per sample per experiment) and the bars are the median and interquartile range. **(D)** Scatter plots of the data from Figure 2D with each experimental repeat plotted side by side. **(E)** γ-ray calibration curve for BrdU alkaline comet assays in DT40. Wild type DT40 cells were incubated with media containing BrdU (100 μM) for 20 h to fully label genomic DNA and then treated with the indicated dose of γ-rays. Alkaline comet tail moments of anti-BrdU stained DNA were quantified under the same electrophoretic and experimental conditions as those applied in BrdU pulse-labelled cells in Figure 2C. The average (±SD) of the median comet tail moments from 2 independent experiments (50 cells per sample per experiment) are plotted. The average fragment size of single-stranded DNA following the indicated γ-ray dose is indicated, calculated on the assumption that each Gy induces ~1100 total DNA breaks (~1000 SSBs & ~50 DSBs)^43–45^ per diploid human genome (12×10^9^ nucleotides) and thus ~1 break every 1×10^7^ nucleotides. Notice that the BrdU alkaline tail moments are insensitive to average fragment sizes below 500 kb, perhaps because fragments below this size are ‘lost’ from the assay during alkaline lysis and/or electrophoresis.

**Extended Data Figure 3.**
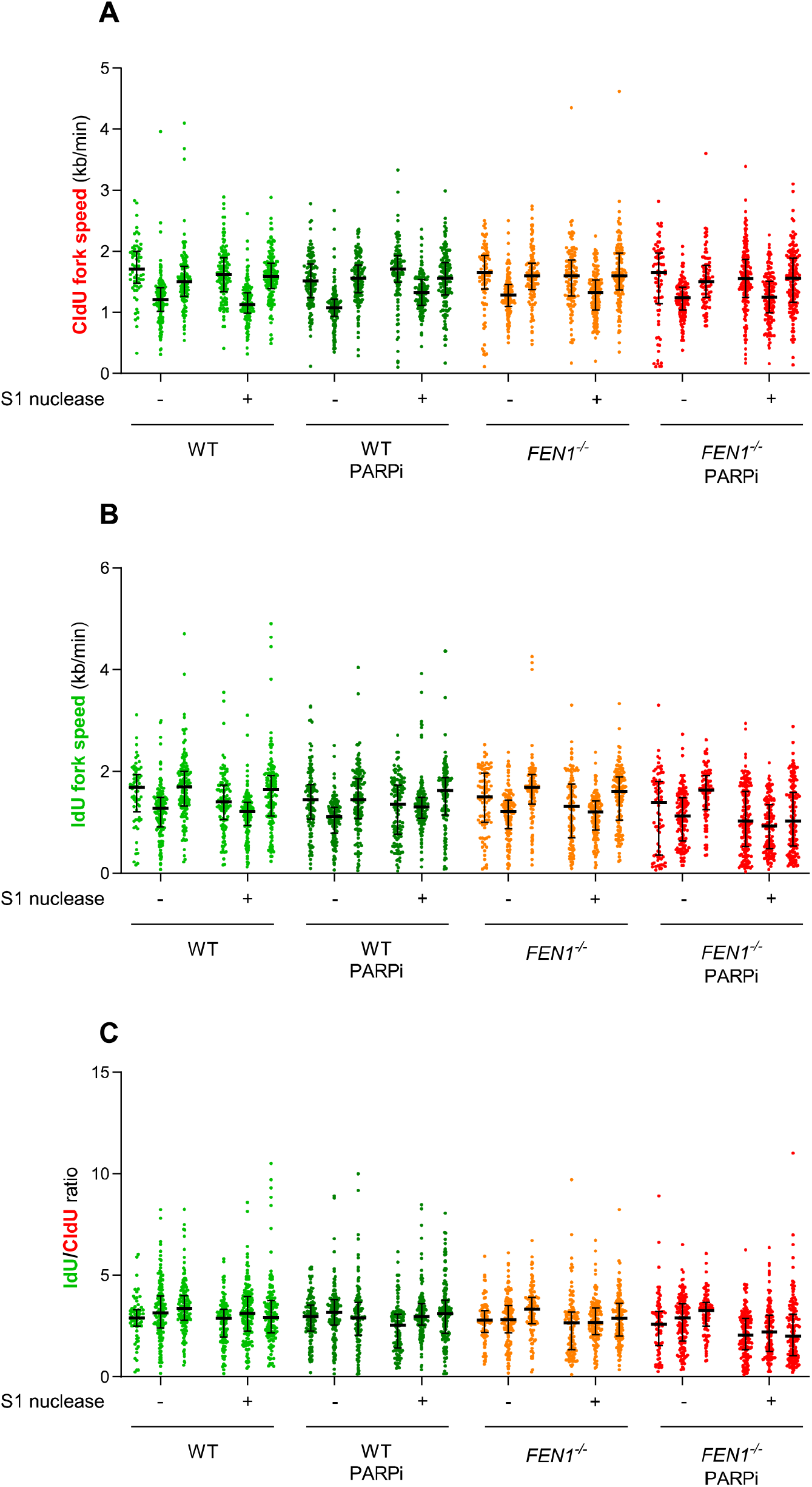
PARP inhibitor induces post-replicative single-strand gaps in *FEN1^−/−^* DT40 cells. **(A-C)** Scatter plots of the data described in Figure 4B-D, respectively, with each data point plotted separately and experimental repeats plotted side by side.

**Extended Data Figure 4.**
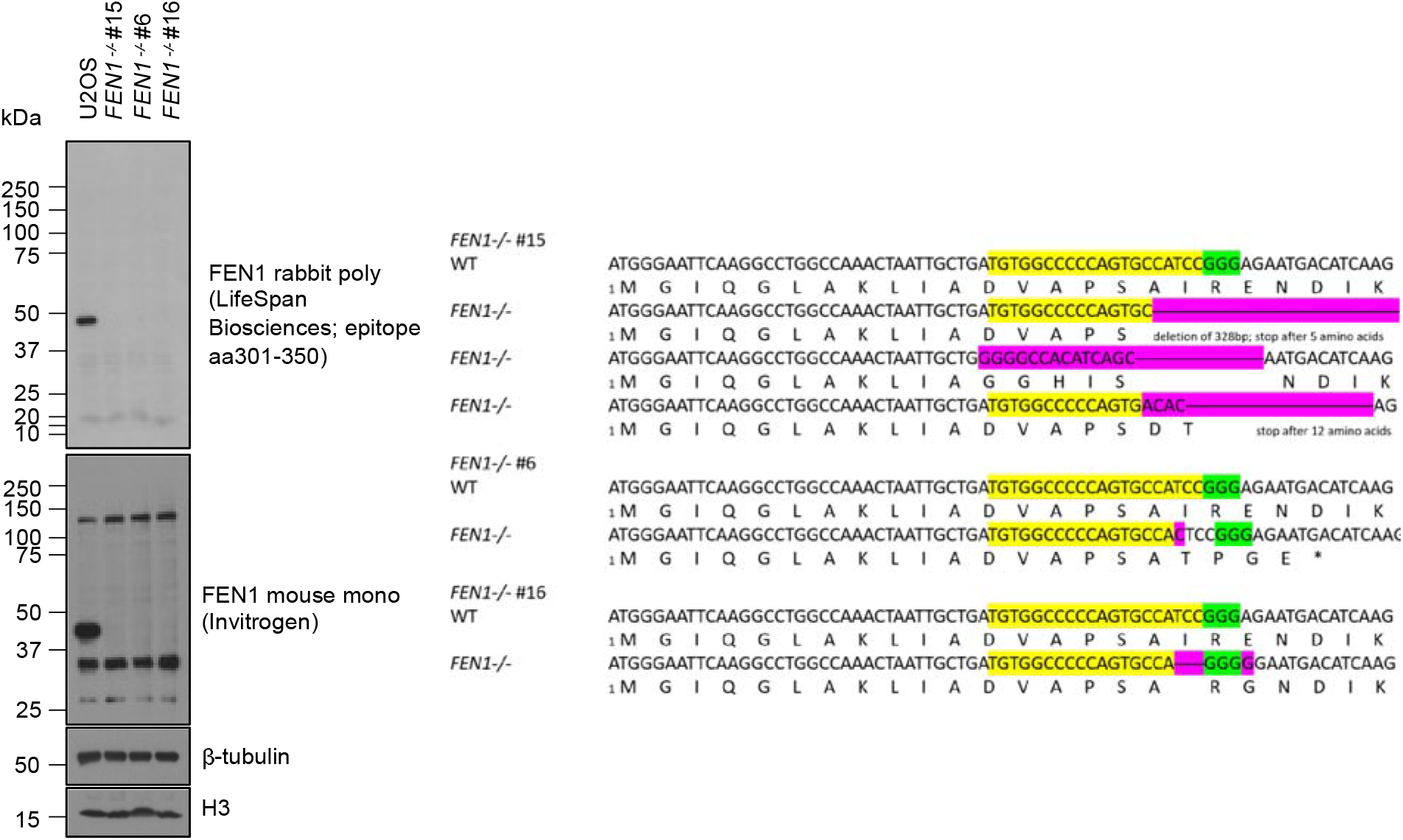
Generation of *FEN1^−/−^* U2OS cells by CRISPR-Cas9 gene editing. **(A)** Analysis of wild type U2OS cells and the indicated *FEN1^−/−^* clones by western blotting of whole cell extracts (*left*) and by Sanger sequencing (*right*) of PCR products of genomic DNA spanning the predicted Cas9 break site. The PAM is shown in green, gRNA targeted sequence is shown in yellow and mutated nucleotides are shown in pink.

**Extended Data Figure 5.**
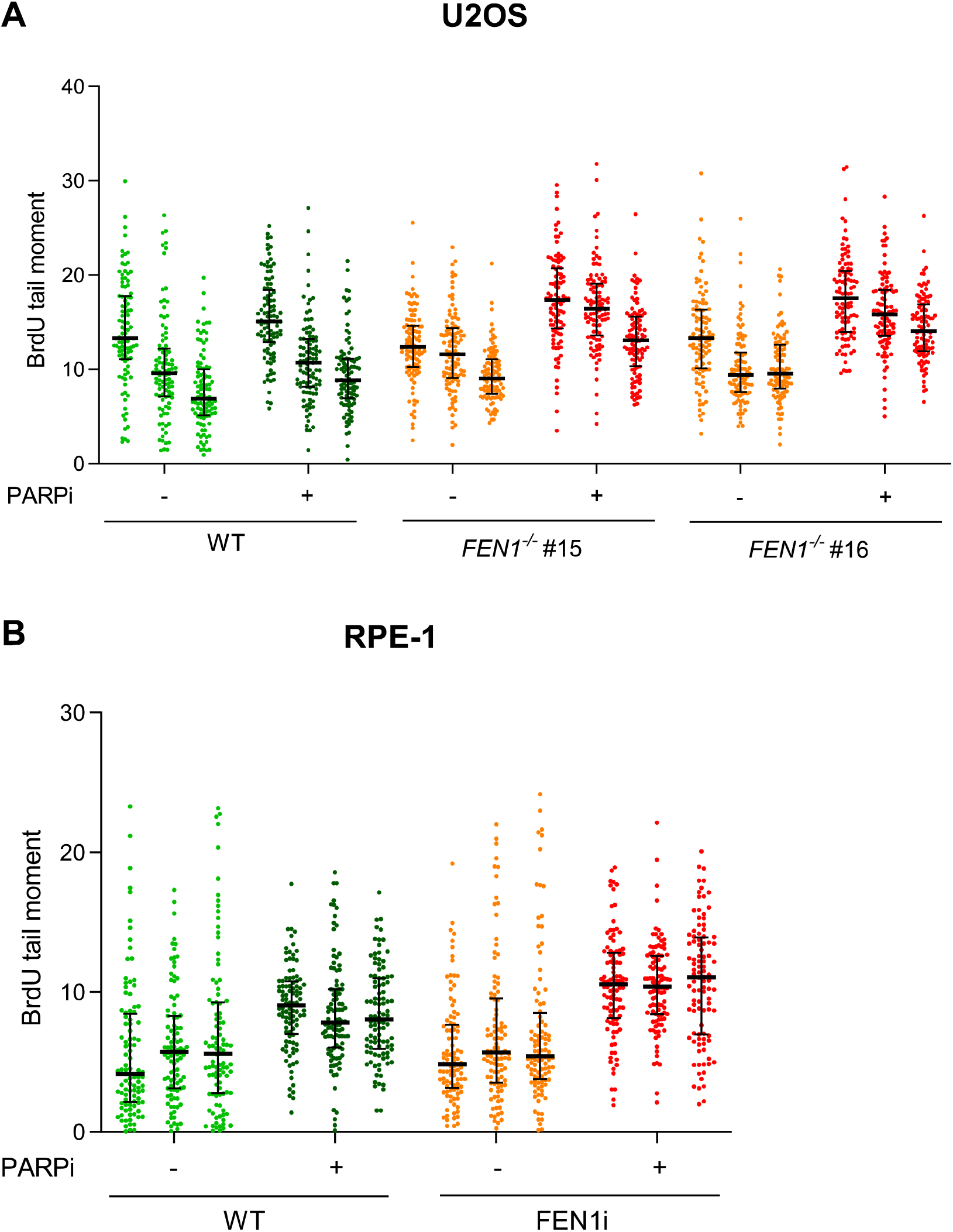
**(A, B)** Scatter plots of the data described in Figure 6A & 6B, respectively, with each data point plotted separately and experimental repeats plotted side by side.

